# Object Colours, Material Properties and Animal Signals

**DOI:** 10.1101/601625

**Authors:** Lucas Wilkins, Daniel Osorio

**Affiliations:** Department of Zoology, University of Oxford, Oxford, United Kingdom; School of Life Sciences, University of Sussex, Brighton, United Kingdom

## Abstract

Colour is commonly regarded as an absolute measure of object properties, but most work on visual communication signals is concerned with colour differences, typically scaled by just noticeable differences (JNDs). Object colour solids represent the colour gamut of reflective materials for an eye. The geometry of colour solids reveals general relationships between colours and object properties which can explain why certain colours are significant to animals and evolve as signals. We define a measure of colour vividness, such that points on the surface are maximally vivid and the ‘grey’ centre is minimally vivid. We show that a vivid colour for one animal is likely to vivid for others, and highly vivid colours are less easily mimicked than less vivid colours. Further, vivid colours such as black, white, red, blue and light, unsaturated shades are produced pure or orderly materials. This kind of material needs to created and maintained against entropic processes that would otherwise degrade or destroy them. Vivid coloration is therefore indicative of ecological affordance or biological function, so that it is valuable to have attentional biases towards these colours regardless of any specific significance.

## 1 Introduction

Animal and plant communication signals carry varied messages: some are attractive while others are defensive; courtship displays appeal to naïve viewers, whereas flowers and aposematic signals need to be memorable. Despite their various functions, and the diversity of colour vision in their natural receivers, signals generally include a limited palette colours including, black, white, saturated hues and light but unsaturated colours such as pink, whereas greys and browns are infrequent.

Any signal must attract attention and engender a response – it will be ineffective if it is overlooked or ignored. Why then should certain colours attract the attention of an eye with any given set of spectral photoreceptors, and why are the same types of spectra appropriate for animals with different types of colour vison? We show here how answers to these questions might be found if a colour is regarded not simply as a measure of the spectral composition of a light, but rather as a property of physical objects. This requires quantifying colour in new ways and dropping two of the most widespread procedures: the splitting of colour into *independent* chomatic and achromatic components, and, the scaling of colour distances using discriminability.

Most work biological signaling treats animal colour vision as means to discriminate between spectra, and is primarily concerned with the magnitudes of colour differences as measured by ‘just noticeable differences’ (JNDs) [Kelber et al., 2003, Kemp et al., 2015, Olsson et al., 2017]. Accordingly, one might predict that a colour signal will attract a receiver’s attention when it differs strongly from the background [Gittleman and Harvey, 1980], or the pattern itself has a high contrast [Rowe and Guilford, 1996, Aronsson and Gamberale-Stille, 2008]. In its everyday use ‘colour’ is not a relative or relational term like ‘contrast’, but is absolute, and part and parcel of object recognition. We do not say “the tennis ball is more yellow than the court,” but “the tennis ball is yellow”. Light reflected from a surface depends upon its chemical composition and (nanoscale) physical structure. Colour vision yields information about these properties. We speak of red faces, blue tea-mugs and so forth, and other animals may be similar. It follows that animals could find particular colours significant because they are characteristic of particular kinds of object.

The significance of a colour might be related to the specific coloration mechanism – for example if a pigment is costly to produce [Olson and Owens, 1998] – or to associations with particular beneficial or harmful objects [Endler and McLellan, 1988, Endler and Basolo, 1998, Palmer and Schloss, 2010]. In contrast with, and complementary to these approaches, we look here more broadly at the relationship between the composition of materials, their reflectances, and their colours. Perhaps surprisingly, we find that there are aspects of object colour that are consistent between different observers, which can be linked to underlying physical properties that are relevent to the psychology of an organism.

In particular, we analyse a property of colour we term ‘vividness’. Highly vivid colours for one organism are highly vivid for any other with the same or more number of photoreceptor types (rules 1 and 2). We then show that highly vivid colours are more informative, in that they correspond to fewer materials, and that those materials will be purer and more ordered than less vivid materials. From this we argue that psychological salience of these colours would be evolutionarily adaptive.

### 1.1 Object Colour Solids and Vividness

Objects are seen by reflected light, and the gamut of all possible reflection spectra can be represented by their locations within in a Cartesian space known as an object colour solid whose axes correspond to photoreceptor excitations relative to the illumination spectrum (Figs 1, 2 Vorobyev [2003], Koenderink [2010]). Colour solids include the three main aspects of colour namely hue, saturation and brightness, whereas the more familiar chromaticity diagrams, such as Maxwell’s triangle discount brightness. An additional difference lies in the nature of the gamut boundaries. In a chromaticity diagram the boundary is defined by monochromatic spectra and (for trichromats) the purple line. As a monochromatic reflection contains negligible light highly saturated (or pure) colours are dark. By comparison object colour solid boundaries include both black (zero reflection) at the origin, and white corresponding to maximal reflection at all visible wavelengths, with intermediate the boundary surface being well approximated by spectral step-functions (Fig 2). Such spectra can be bright, and are more nearly physically realizable by natural pigmentation.

**Figure 1:**
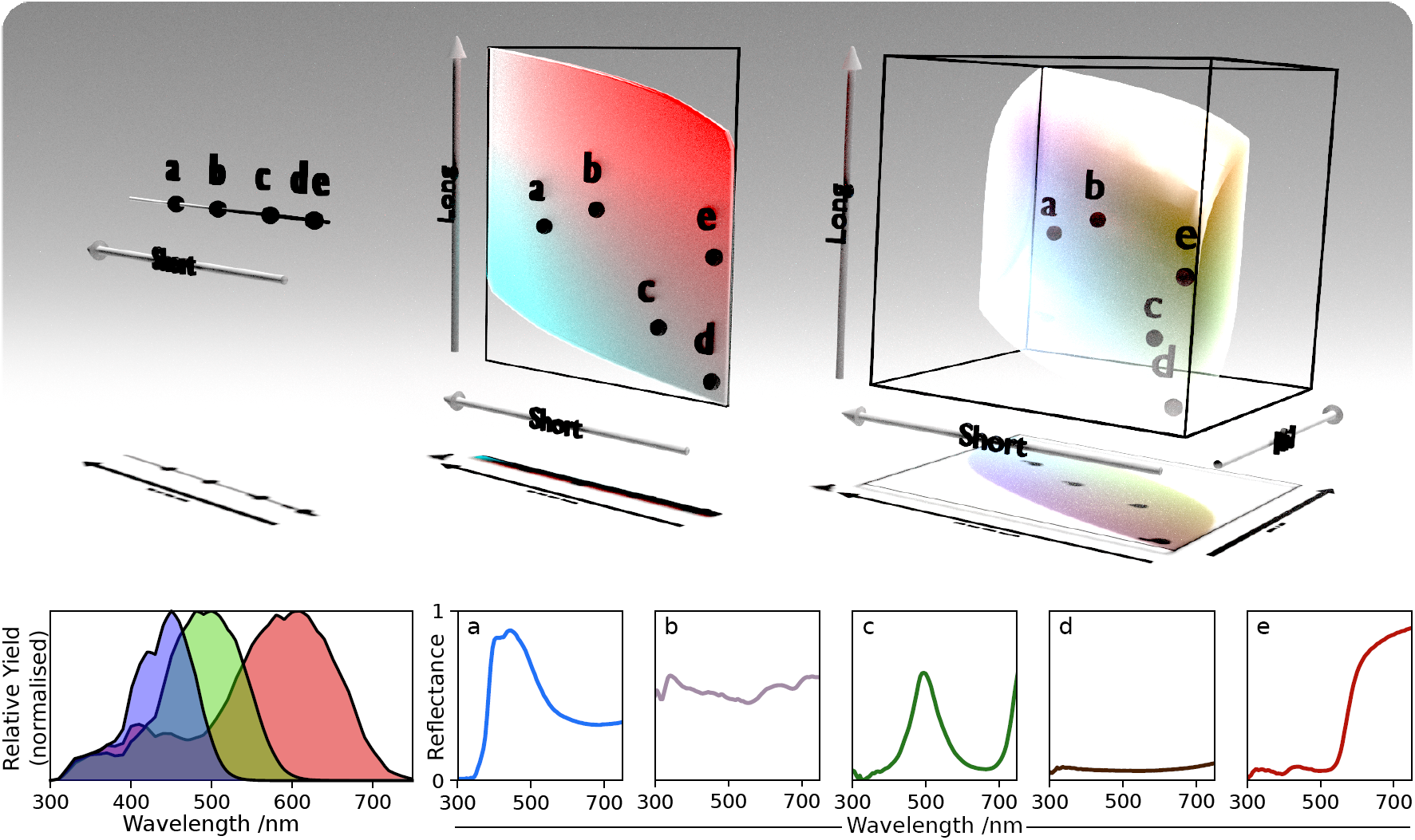
The Object Colour Solid *Top, left to right:* Colour solids for a monochromat, a dicromat and trichromat. *Bottom left:* Relative quantum yield curves (spectral sensitivity multiplied by the illuminant and normalised). The monochromat has just the short wavelength sensitive photoreceptor class, the dichromat contains the short, and the long wavelength sensitive photoreceptor classes, the trichromat contains all of them. Each ‘expands’ the solid outwards into the new dimension. Note how the positions of colours (a-e) in the pre-existing dimensions do not change as more photoreceptor classes are added. *Bottom right:* The natural reflectance spectra responsible for colours (a-e).

**Figure 2:**
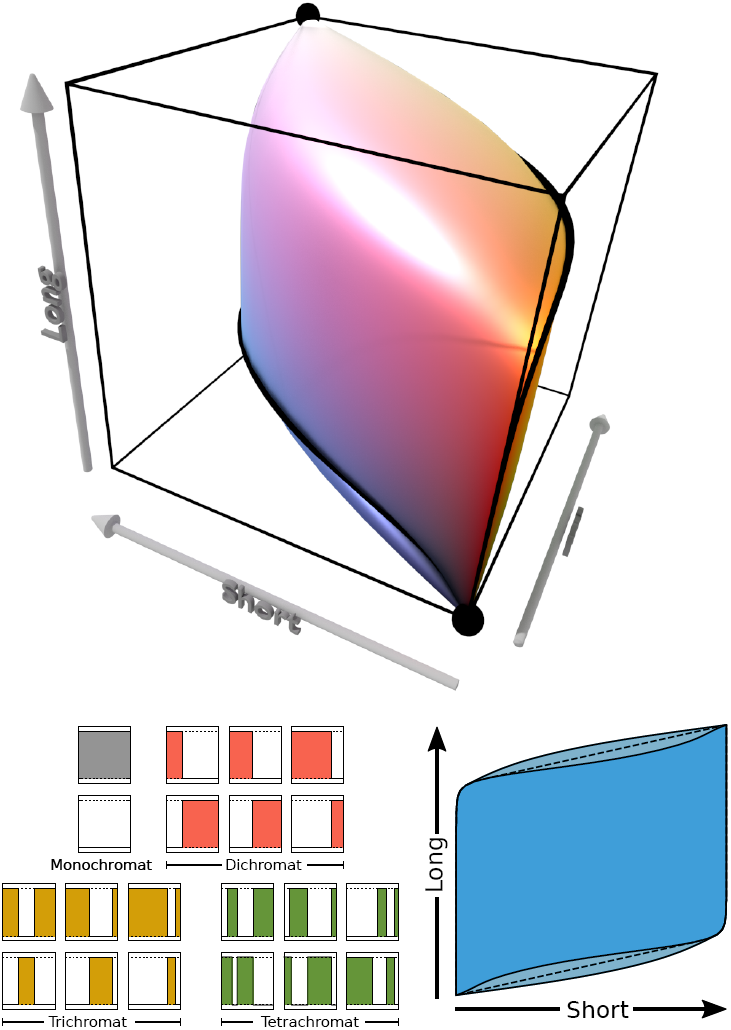
The Schrödinger approximation. *Top:* Loci of Schrödinger spectra for the trichromat in figure 1. The coloured surface shows the trichromatic Schrödinger spectra. Note that for this observer the surface formed by Schrödinger spectra is concave in some places (see main text), this is most easily observed in the blue region. The black ribbon running around the edge of the solid shows the dichromat Schrödinger spectra. The spheres at the tips correspond to the monochromat Schrödinger spectra (i.e. back and white). *Bottom left:* Examples of Schrödinger spectra for monochromats, dicromats and trichromats. The relationship between the colour solid and chromaticity (colour triangle) has been explored by others [Wyszecki and Stiles, 2000, Luther, 1927, Nyberg, 1928]. *Bottom right:* A dicromatic solid (same as in figure 4) showing the locus of the dicromat Schrödinger spectra (innermost shape). The non-convexity is much easier to see in this rep-resentation. The convex hull of the Schrödinger spectra’s colours is shown as a dotted line, this does not reach the boundary in its entirety either. For this observer the dicromatic Schrödinger spectra have a mean vividness of ≈ 0.943 (all in the range ≈ 0.843 to 1). The colours used to render these solids are only indicative of their true appearance.

Following the logic that colour refers primarily to the physical properties of reflective materials (or objects) we propose a measure of colour within the object colour solid which we call vividness. Vividness resembles colorimetric parameters such as purity or saturation, but achromatic colours – black and white – and light unsaturated colours can be highly vivid. We show mathematically and empirically that the vividness of reflectance spectra is well correlated between different types of colour vision. Consideration of the relationship between vividness and the physical processes that generate object-colour demonstrates that orderly nanostructures or pure pigments are typically more vivid than their less orderly or pure materials. As order emerges against entropic tendencies, vivid or ‘bright’[Hamilton and Zuk, 1982] coloration is indicative of a functional role, and therefore more “meaningful”, and hence such colours worthy of greater attention.

## 2 Modelling

We start with an account of why the achromatic colours black and white are maximally vivid, leading to a general model which includes chromatic colours.

### 2.1 Black and White

Models of colour vision and colour appearance usually treat chromaticity, which combines hue and saturation, as qualitatively distinct from brightness or luminance. This distinction is grounded in physiology and psychophysics [Livingstone and Hubel, 1988, Osorio and Vorobyev, 2005], yet black and white are ‘colours’ in ordinary English usage, and they are common in biological signals. In physiological terms an animal sees black when its spectral photoreceptors all have a low excitation, and white when they all have a high excitation. Hence, spectra that look black have low intensity at all visible wavelengths and spectra that look white have high intensity. As there is only one perfectly black spectrum, and a perfectly white surface must reflect maximally at all wavelengths it follows that spectra responsible for black and white are the same for all observers that share the same range of visible wavelengths, with minimal or maximal excitation across that range. These spectra can be designated as ‘extreme’, because they are at the limits of receptor excitation achievable by a surface.

Unlike black and white, intermediate reflectance spectra can produce different excitations according to the particular set of photoreceptors in a given eye. For the simplest case of two eyes each with a single type of photoreceptor, but tuned to different wavelengths, a stimulus of intermediate intensity can give different receptor responses in the two monochromatic observers. Consequently the appearance of greys for different types of monochromat is less predictable than are black and white. Similarly, for any given monochromatic eye only one spectrum can give black or white, but many different spectra can produce indistinguishable intermediate responses, a phenomenon which is known as colour metamerism[Logvinenko, 2009].

Black and white, produced by extreme spectra, are maximally vivid colours, while have greys have lower vividness.

The best known geometric representations of colour are chromaticity diagrams in which desaturated colours lie near the centre and the most saturated colours at the extremities, with the angle around the centre specifying hue.

One such diagram is Maxwell’s triangle [Maxwell, 1860]. Using a method based on colour mixing, Maxwell specified the colours of monochromatic lights as mixtures of red, green and blue primaries. When the brightness is ignored these results can be drawn in a triangle, with pure primaries at each corner, and mixtures within. The line of all the monochromatic lights, the “monochromatic locus” or “spectral line” forms a rounded Λ shape along two of the edges, the ends joined by a “purple line”. The fractional distance from the centre to the edge is then a measure of spectral purity, which is related to the colour’s perceived saturation [Wyszecki and Stiles, 2000].

For colour vision of non-human species similar diagrams are typically based on photoreceptor spectral sensitivities[Renoult et al., 2017]. For an eye with *n* spectral types of photoreceptor (contributing independently to colour vision) the chromaticity diagram is *n* − 1 dimensional. Chromatic spaces are good for describing the colours of lights, but less suitable for reflectance spectra, because a *reflectance* can only approach the boundary by reducing the amount of reflected light, making the colour darker – maximal spectral purity is black! Hence chromaticity diagrams typically exaggerate differences between dark colours.

Object colour solids are so-named because they appropriate for representing ‘object’ or reflectance spectra. They are useful in colour reproduction and the formulation of dyes and pigments, because the available gamut can be compared to the colour range visible to the human eye. The axes of colour solids [Wyszecki and Stiles, 2000, Schrödinger, 1920, Vorobyev, 2003, Koenderink, 2010] correspond to photoreceptor excitations (or a similar set of primaries) normalised to the illumination intensity. As photoreceptor spectral sensitivities overlap, colour solids do not fill the space defined by the axes, but are roughly ellipsoidal with two pointed corners (Fig 1). Monochromatic spectra lie an infinitesimal distance from the origin which is black, and maximal reflectance (white) is at the opposite vertex. Humans have three types of cone photoreceptor, and hence a 3-dimensional object colour space, but the same geometrical principles apply to any type of colour vision: for most mammals, which are dichromats, the space is 2-dimensional, whereas the spaces of birds are probably 4-dimensional [Kelber et al., 2003, Vorobyev, 2003]. As there are both empirical and mathematical relationships between these spaces we refer to all of them as colour solids.

### 2.2 Colour Solids and Vividness

We now more formally define the colour solid and vividness. For a given set of photoreceptor spectral sensitivities (*s*_*i*_(*λ*), *i* ∈ 1 … *n*) and an illuminant (*l*(*λ*)), reflectance spectra can be organised into an geometric object known as the *object colour solid* [Schrödinger, 1920, Wyszecki and Stiles, 2000], shown in figure 1. This object is formed from the colours (a vector of photoreceptor quantum yields, (*q*_1_ … *q*_*n*_)) associated with all theoretically possible reflectance spectra: that is all distributions of reflectance values between zero and one, taken over visible wavelengths (Λ):

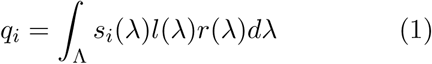

To obtain the colour object solid, photoreceptor responses (*q*_*i*_) are normalised to the quantum yield of a perfectly reflecting surface^1^ 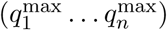.

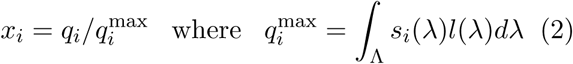

This means that black has *x* coordinates of (0, 0, *…*) and white has coordinates (1, 1, *…*). Throughout we express the *x* coordinates in terms of a relative quantum yield function 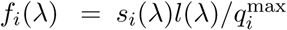 :

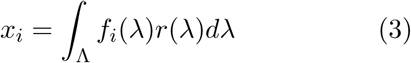

The object colour solid is convex, and lies within a unit *n*-cube (Fig 1). It is pointed at the diagonally opposite black and white corners, but the other corners and edges are smoothed due to the spectral overlap of the photoreceptors. Except for the boundaries, every point in the solid maps to more than one reflectance spectrum, corresponding to colour metamerism[Logvinenko, 2009]. The boundaries represent reflectances with the highest spectral purity for a given luminance. Unlike chromaticity diagrams, the colour solid therefore accounts for the trade-off of saturation against luminance, so light colours can lie on the boundary. This accords with the intuition that we do not see light colours as necessarily less pure than dark.

The exact calculation of the colour solid is rather involved. Numerical solutions can be obtained by dynamic programming [Wyszecki and Stiles, 2000] or our own method (see SI 2 for details of both), but for the current purpose it is better to start with Schrödinger’s approximation [Schrödinger, 1920, Vorobyev, 2003], which uses spectra formed by step changes in intensity between zero and one. Figure 2 illustrates Schrödinger’s spectra. The number of step changes depends on the number of spectral classes of photoreceptor: for an eye with *n* spectral classes the maximum number of step changes needed to approximate the colour solid boundary is *n* − 1. For dichromats, they are single steps, the two series (step up and step down) ranging from black through red, orange, then yellow to white, and from black through blue then cyan to white. For trichromats there are two steps in the visual range, while for a tetrachromat the Schrödinger spectra include those that have a reflectance of 0 up to some wavelength *λ*_1_ then 1 until another, *λ*_2_, then 0 until *λ*_3_ then 1, and in addition their inversions, i.e. those that have values of 1 then 0 then 1 then 0. Whilst the Schrödinger spectra are only approximations to the extreme spectra that lie on the boundary of the solid, the extreme spectra, like Schrödinger spectra, will only ever have either maximal (1) and minimal (0) intensity at every wavelength^2^

Schrödinger’s approximation of boundaries of the object colour solid holds best when photoreceptor spectral sensitivities have a single peak. Some receptor spectral sensitivities are bimodal (e.g. Fig 3), in which case Schrödinger’s spectra typically lie slightly inside the boundaries of the corresponding colour solids. Such deviations are in practice small, because photoreceptor spectral sensitivities are typically smooth with a dominant peak. As long as this approximation holds a conclusion analogous to that for black and white applies; namely that for eyes with the same number of photoreceptor classes, the Schrödinger spectra corresponding to boundary colours are the same. Differences between receptor sensitivities mean that the shape of the object colour solid varies between species, and the locations of individual spectra within the boundary are not directly comparable (Fig 3), but they will be on the boundary nonetheless. It follows that Schrödinger spectra for one eye are Schrödinger spectra for an eye with a larger number of photoreceptor classes: the set of Schrödinger spectra for an *n*-chromat contains the Schrödinger spectra for an *m*-chromat, when *m* ≤ *n*.

**Figure 3:**
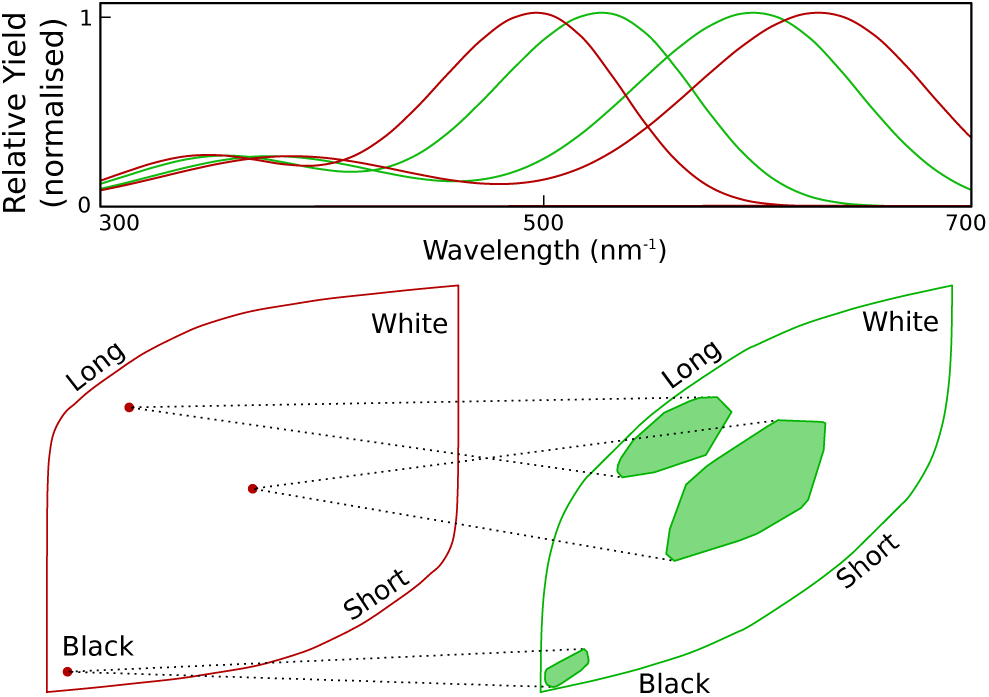
The relationship between the colours in two dichromat object colour solids. Because of metamerism, points on the left correspond to multiple points (areas) on the right. The exact location depends on the particular underlying spectrum. Similarly (though not shown) points on the right correspond to areas on the left. At the edge of the solid the correspondence is one-to-one, and the size of the region in one corresponding to a point in the other increases towards the centre. These mapped areas, like the solids themselves, are based on the assumption that spectra may instantaneously transition between zero and one. Such transitions cannot be realized physically, and the spaces and areas are theoretical bounds. The filled areas on the right are bounded by spectra with a greater number of transitions than the solid. These filled areas are conservatively large estimates of the degree of metamerisim that is physically realisable. The relative quantum yields were produced from an A1 pigment template which has a pronounced *β* peak [Govardovskii et al., 2000], but the correlation in vividness remains even though the Schrödinger spectrum approximation does not hold exactly for these double-peaked relative quantum yield functions.

The tendency for the same spectra to lie at the boundary of the colour solid extends to colours not on the boundary, although, as was the case for black, white and grey, the variablility in the position is greater for more central colours (Fig 3). Whilst the theoretical limitations on the position, illustrated in Fig 3, are broad, the spectra that achieve these limits are difficult to realise practically. This is evident in Fig. 4.

**Figure 4:**
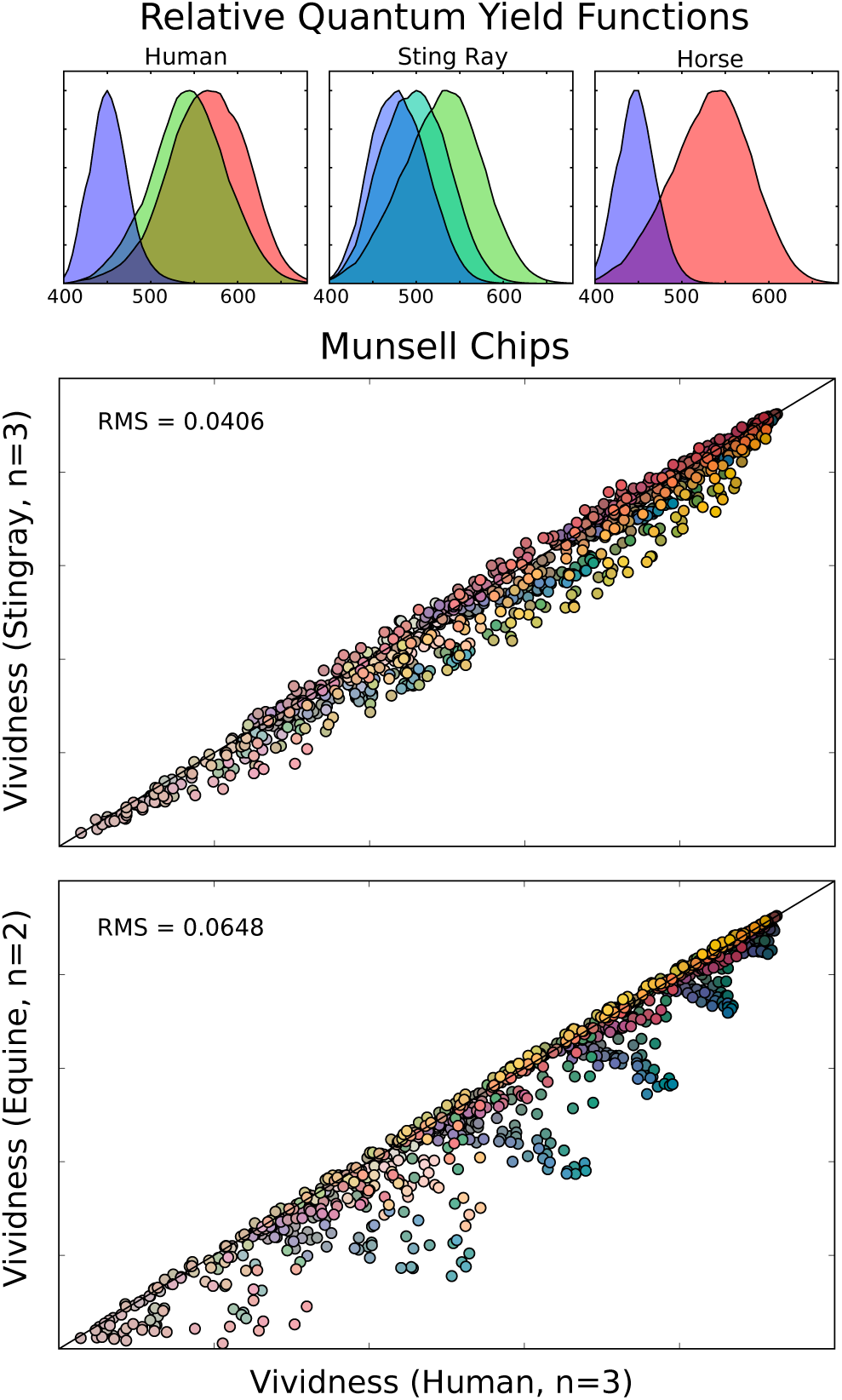
Demonstration of the correlation of vividness between three disparate species. The *top row* shows example relative quantum yield functions with D65 illumination Wyszecki and Stiles [2000] for trichromatic humans Stockman and Sharpe [2000] and sting rays, Hart et al. [2004] and dichromatic horses Carroll et al. [2001]. The scatter plots compare the vividness of the Munsell chips [Munsell et al., 1950, Parkkinen et al., 1989] – a collection of coloured stimuli designed to cover the human colour gamut uniformly – for the three species. The *middle* plot demonstrates *rule 1* – the approximate equality between vividness for the two species with the same number of cone classes (humans and sting rays). The *bottom* plot shows, in addition, the effects of *rule 2* – whist the approximate equality holds for many cases, some chips which we see as blue or pink can be less vivid for a horse.

In practice, spectra of natural objects are a small subset of physically possible spectra [Maloney, 1986, Osorio and Bossomaier, 1992, Vorobyev et al., 1997], and natural spectra tend to be smoother than Schrödinger spectra [Maloney, 1986] (compare Fig 1 and Fig 2). Also, spectra with multiple transitions within the visible range are unusual, so that higher order Schrödinger spectra are seldom approached in nature; exceptions include some structural colours, such as that found on the nape of the feral pigeon, which can appear greyish to trichromats, but is likely to be vivid for birds [Osorio and Ham, 2002].

### 2.3 Definition of Vividness

We define vividness as a number ranging from zero at the centre of the colour solid to one at the boundary, which is linear with respect to the colour solid coordinates (*x*_*i*_). For an observer/illumination combination that is described by *n* relative quantum yield functions (*f*_1_ … *f*_*n*_) the vividness of a reflectance *r* is:

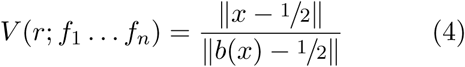

where *b*(*x*) is the position of the boundary in direction of the vector *x* − ½ from the centre (½).

The numerator is the Euclidean distance from the centre of the solid. The division by ‖*b*(*x*) − ½‖ maintains an invariance between species. It has the effect of adjusting the numerator to reflect more physical, rather than perceptual, properties. The latter being more difficult to assess for non-human species (and even for humans).

### 2.4 Properties of Vividness

The properties of colour solids we have discussed so far can be formally expressed with two rules, the first is motivated by our argument above and it is true insofar as it is approximate, and contingent on having “well behaved” relative quantum yield functions, the second is a geometric fact:

#### Rule 1: Approximate Equality

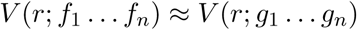

For two observers with equal numbers of spectral receptor classes (*n*), the vividness of a stimulus is approximately the same for both observers. This rule is motivated by the foregoing discussions, and is corroborated by figures 3 and 4. For more vivid colours the range of discrepancy decreases, and the approximation is better – a consequence of there being “fewer” metamers. This phenomenon can be seen in figure 3.

As this rule concerns the relative quantum yield functions *f*, which are obtained from both spectral sensitivities and the illuminant, rule 1 is both a statement about changes in photoreceptor sensitivities and in illumination.

#### Rule 2: Monotonicity with Dimensionality

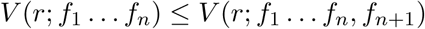

For a given illumination, an increase in the number of receptor types will result in an increase in vividness, so that colours are more vivid for species that have a large number of photoreceptor classes (e.g. birds) than for those with fewer (e.g. mammals). This effect can be observed in figure 4, and a proof is given in SI 1.

These rules^3^ allow one to describe the relationship between any two observers.

As vividness increases the constraints on the variety of spectra that can realize the colour become increasingly restrictive, until, at the boundary of the solid there is a unique spectrum. As there are “more”^4^ spectra that correspond to each of them, less vivid colours are more prone to metamerism. This rule can be compared to the metamer mismatching transformation that occurs under variable illumination [Tokunaga and Logvinenko, 2010].

Colour *purity* [Wyszecki and Stiles, 2000] resembles vividness for the chromatic plane, and has fairly similar properties. This is because the chromaticity space is a scaled cross section of the colour solid at fixed luminance as it goes to zero (i.e. a slice of the solid very near black). Vividness also resembles other measures, such as *chroma* [Endler, 1990]. In addition to the difficulties we have already highlighted with chromaticity, purity is not defined for monochromats, is not mathematically well behaved for dichromats, and can be quite complex for tetrachromats and above (see supplementary material SI 3).

### 2.5 Mixing

Vividness has a fundamental relationship to physical order, as exemplified by the case of conservative mixing. That is to say, where two colours, *x*_*a*_ and *x*_*b*_, are mixed in a ratio of *z*_*a*_ : *z*_*b*_ and the resulting colour is:

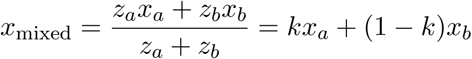

where *k* = *z*_*a*_*/*(*z*_*a*_ + *z*_*b*_) and the various values of *k* correspond to various positions on the line segment between *x*_*a*_ and *x*_*b*_. This holds regardless of the spectra that produce *x*_*a*_ and *x*_*b*_.

The distance of points on this line segment are closer to (or the same distance from) the centre of the solid than the more distant of *x*_*a*_ and *x*_*b*_. If the solid were a perfect sphere (i.e. if the distance to the boundary were fixed) we could conclude directly that vividness of the mixture was smaller than that of the colours being mixed. As the solid is not a sphere, we must make a further observation: where the surface is (strictly) convex, a line segment connecting two points on the surface of the solid passes entirely within its volume (for a proof see supplementary material SI 1). The colour solid’s boundary colours have the distinguishing feature that they cannot be made by a mixing other object colours.

The rule applies to all observers, in spite of the fact that numerical values of *V* for a particular object colour may well be different.

#### Rule 3: Convexity of Mixing

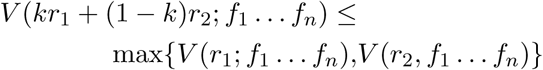

for all *k* ∈ [0, 1]. Mixing two colours results in a colour that is less vivid than the most vivid of the two (and perhaps less than both). Rule 3 can be written in a more general form, for a mixture of multiple colours, as:

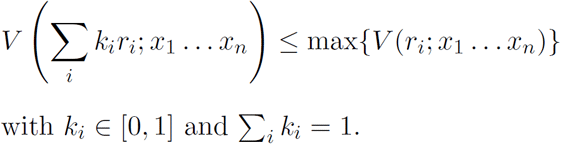

This expression describes a phenomenon familiar to anyone who has mixed paints, once a duller colour is mixed into a more vivid one there is no way to recover the original vividness except by adding an even more vivid paint.

This is not a property unique to vividness, but we must bear in mind the significance of this rule for our argument: Without this rule we would only have mathematical results about the geometry of colour solids, but with this rule, we can talk about the physical properties of vividly coloured materials, and thereby talk about their ecological function and how they are perceived (in a Gibsonian sense [**?**], at least).

## 3 Discussion

Object colour solids are useful representations of the colours of reflective surfaces, as opposed to lights. The mathematical properties of these spaces along with empirical evidence (Fig 4) leads to three main findings relevant to the evolution of colour in communication signals.

Firstly, we define a measure of colour, vividness, which corresponds to the distance of a colour from the (grey) centre of the soild, and show mathematically that a spectrum which is highly vivid (i.e. near the boundary of the colour solid) for one type of colour vision will be highly vivid for any type that has the same number or more spectral types of photoreceptor. We find empirically that the vividness of natural reflectance spectra is correlated for eyes with different sets of photoreceptors (Figs 4 and 3).

Secondly, colours on the boundary of the solid are attributable to a single reflectance spectrum, and the number of spectra that map to a given point in the colour solid increases towards the centre, so that many spectra can look ‘mid-grey’. Highly vivid colours are therefore more likely to be associated with a specific physical cause (i.e. material), and they will be less easily reproduced by alternative means than less vivid colours. Thirdly, mixing colours inevitably reduces the vividness of the more vivid colour, so that pure materials tend to be more vivid than mixtures. This is why vacuum-cleaner dust is greyer than home furnishings. For structural colours increasing regularity of nano-structure increases vividness, and mixing of pigments can only render them less vivid.

Due to entropy order does not arise by chance in nature, so vivid colours are indicative of some functional role. This need not be as a signal; the vivid colours of leaves and blood are due to high concentrations of light harvesting and oxygen transport pigments. This is not to say that and a dull coloured tissue cannot have a specific function, the implication only works one way – vividness requires some kind of order, but order does not necessarily result in vivid colour. Nonetheless, if an object has a functional role it is *a priori* worthy of attention, which is a requirement for any signal.

Historically, vivid coloration has been associated with the phenonemon of life, and this in turn has been related to thermodynamics. In *Tropical Nature* [Wallace, 1878] Wallace argues that colourfulness is a consequence of “vital energy”, and hence a natural attribute of living organisms. While Schrödinger proposed in his book *What is Life?* [Schrödinger, 1944], that life is characterised by order away from chemical equilibrium, that is by being non-entropic. Here we have seen why vivid colours are indeed likely to be associated with a system – such as life – that counters the effects of entropy.

### 3.0.1 Aposematism and Mimicry

The strong contrasts and vibrant colours of aposematic displays illustrate some of our findings. Vivid colours are salient, so they have the potential to promote both innate and learnt responses, and they and will be seen consistently by a broad range viewers, so they need not be predator-specific. For instance in tropical forests predators of insects include monochromatic strepsirrhine primates, dichromatic mammals and snakes, trichromatic primates and amphibians, and tetra-chromatic birds and lizards [Kelber et al., 2003]. A consequence of rule 2 is that the achromatic extreme colours – black and white – will be effective for all receivers, those approximating the dichromatic Schrödinger spectra (which we would see as two series black, red, yellow, white and black, blue, cyan, white) would be effective for dichromats and above, the trichromatic series which adds purples and greens for trichromats and above, and so on. Similar principles apply in marine environments where predator colour vision ranges from monochromacy in cephalopods through di, tri and tetrachromacy in various fish to the multispectral system of stomatopods[Marshall et al., 2015].

A further benefit of vivid defensive signals arises because as vividness increases copying a colour entails more exact matching of the spectrum. A defended aposematic model may in principle outmanoeuvre a Batesian mimic by adopting more vivid colours outside the mimic’s physiological ‘gamut’ [Franks et al., 2009, Briscoe et al., 2010, Bybee et al., 2011] and conversely there might be a pressure for Müllerian mimics to share less vivid colours.

### 3.1 Modelling of Colour in Biological Signals

Investigation of colour signaling by animals and plants often starts by modelling photoreceptor responses to reflectance spectra. Receptor responses do not directly specify colour differences or colour appearance, which normally requires recourse to psychophysical models [Kelber et al., 2003, Kemp et al., 2015]. Models based on chromaticity assume that lightness (or luminance) is discounted, and although some add achromatic contrast as a separate parameter[Siddiqi et al., 2004, Olsson et al., 2017], all such models can lead to difficulties. For example they predict that dark colours are unrealistically distinct, and there may be an implicit assumption that the strong achromatic contrasts, which are present in many signals are either irrelevant or have a qualitatively different function from chromatic components.

Object colour solids represent the full gamut of colours visible to an eye, and importantly offer a natural means of representing colour as a property of reflective materials rather than spectral lights. It is straightforward to define the locations of reflectance spectra in the object colour solid, and hence to explore broader questions about the gamuts of colour signals directs at various types of receivers (cf [Osorio and Vorobyev, 2008]. Vividness is a simple and well defined measure of colour, within a colour solid which can be related to the physical properties coloured materials, and gives insight into how and why animals with diverse visual systems might evaluate colour. Of course it remains an empirical question whether vividness is a useful measure of colourfulness. One could test whether vividness should predict attention or salience better than colour saturation or purity, or to a scale based on colour distances measured in terms of just noticeable differences within the colour solid.

## 4 Conclusion

In *The Origin of Species* Darwin writes. “…the belief that organic beings have been created beautiful for the delight of man, … has been pronounced as subversive of my whole theory, …” [Darwin, 1859]. Contemporary literature rarely addresses this question directly, but offers a range of accounts of why certain colours or patterns should be attractive to animals [Zahavi, 1975, Grafen, 1990, Guilford and Dawkins, 1991, Anderson, 1994, Johnstone, 1995, Endler and Basolo, 1998]. Some refer to the nature of the sensation, and postulate general aesthetic principles, whereas others highlight the specific value of a stimulus. For example it is debated whether animals use particular colours in courtship displays because they resemble objects of value such as food items [Allen, 1879, Endler and Basolo, 1998] or because the colours are indicative of the quality of a potential mate[Hill, 1991]. Here we find that these distinctions may be blurred, because consideration of the physical causes of the coloration suggests that certain colours should *a priori* be significant regardless of specific associations with objects of relevance to the animal, or indeed its particular type of colour vision. There is no need to invoke evaluative judgements [Prum, 2012] or co-evolutionary processes [Fisher, 1930] such as those associated with sexual selection. In addition the laws of thermodynamics mean that there is an immediate cost to producing pure materials and hence vividly coloured tissues, but this is not part of the main thrust of our argument, which is instead about a general psychological bias towards vivid colours.

Models of visual salience to humans typically include components that are akin to vividness [Niebur and Koch, 1996], and this kind of low level prediction can be compared and contrasted by higher level (evaluative) theories based on asking subjects about the aesthetic value of a colour. Palmer and Schloss’s [Palmer and Schloss, 2010] valence theory proposes that humans prefer colours associated with desirable objects, to those associated with decay, excrement and so forth. Given that decomposition and biological waste tend to be chemical mixtures, with low vividness, whereas pure materials tend to be more vivid it would be interesting to test whether vividness predicts colour preference as well as the valence. At a more practical level, we can offer some assurance to field biologists that it is reasonable to generalise from our own colour perception to that of other animals, despite their physiological differences [Bennett et al., 1994].

## Data Availability

The code used to generate figures here have been made available, along with documentation and usage examples, as an open source project currently hosted at https://github.com/lucas-wilkins/lemonsauce. The numerical methods are also detailed in the supplementary information.

The Munsell chip data are available from the Spectral Color Research Group, University of Eastern Finland at http://www.uef.fi/web/spectral/-spectral-database [Parkkinen et al., 1989].

## Acknowledgments

We would like to thank Mary Stoddart, Almut Kelber and Justin Marshall for their comments, and Justin Marshall also for the spectra used in figure 1.

## Supplementary Information

### SI 1 Proofs

#### SI 1.1 Convexity and Symmetry

The convexity and symmetry of the colour solid is well established, but it is useful to quickly visit it, as many of the appendices use this fact.

The (weak) convexity of the colour solid means that if colours *x* and *y* belong to it, then so do all the colours lying the line between them, i.e. *kx* + (1 − *k*)*y* for all *k ∈* [0, 1]. Because *x* and *y* are assumed to be colours within the solid, there are corresponding reflectances *r*_*x*_ and *r*_*y*_ bounded between 0 and 1 at all wavelengths.

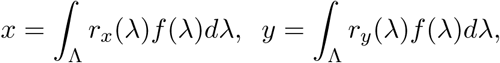

The reflectance *kr*_*x*_(*λ*) +(1 − *k*)*r*_*y*_(*λ*) will result in the colour *kx* + (1 − *k*)*y*, and as this is a convex combination of the reflectance *r*_*x*_ and *r*_*y*_ this will always have values between *r*_*x*_ and *r*_*y*_, and there-fore between 0 and 1. Thus this colour will always belong to the colour solid.

A similar argument applies to the central symmetry of the colour solid. If *x* is a point within the solid, we can find at least one reflectance *r*_*x*_(*λ*) that will generate it (by definition). For every reflectance *r*_*x*_ bounded between 0 and 1 there is a corresponding reflectance 1 – *r*_*x*_(*λ*) which is also bounded between 0 and 1. So, there will be a corresponding colour, 1 − *x*, within the solid. We can also see from this that the centre of the solid 1/2 will be mapped to itself through this operation, and thereby corresponds to the solid’s center of inversion.

#### SI 1.2 Rules 2 and 3

The proofs of rules 2 and 3 can be demonstrated by planar geometry using only the facts that vividness is, by definition, the fractional distance from the centre of the solid to the boundary, and that the colour solid is convex.

The first of these is a matter of definition, and the second follows from the space of reflectances being convex (essentially an infinite dimensional cube) and the integral ∫_Λ_ *r*(*λ*)*f*_*i*_(*λ*)*dλ* acting as a linear projection of *r*.

#### SI 1.2.1 Monotonicity with Dimensionality

The proposition to be shown is:

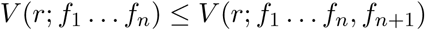

This states that the vividness in a colour solid either increases or stays the same when an extra dimension (spectral photoreceptor class) is added. This may be interpreted using the following diagram

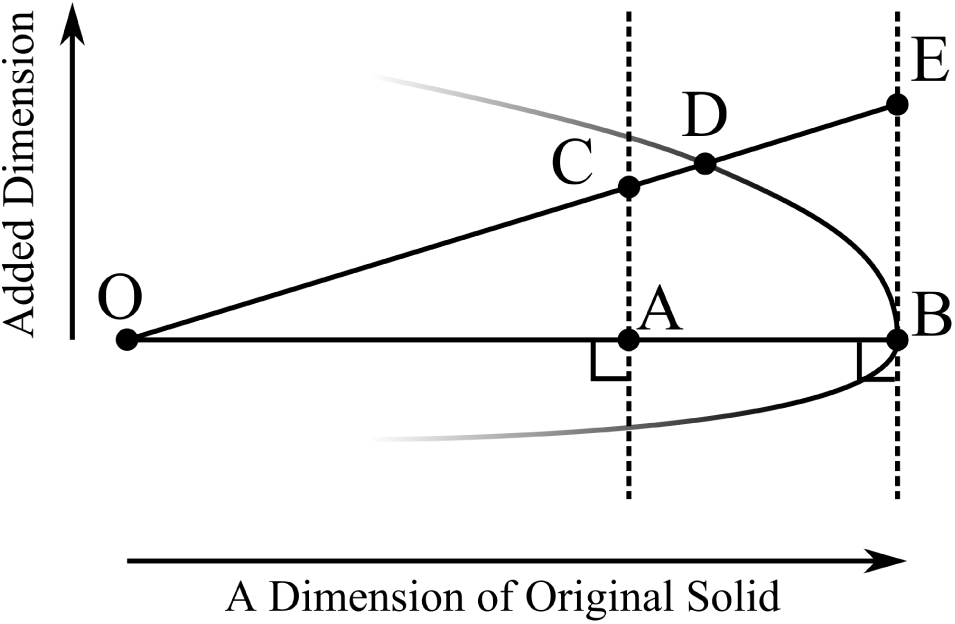

Left to right is the component of the original solid parallel to the vector from the centre of the solid, *O*, to the colour *A*. The corresponding boundary colour is labeled *B*. The upwards direction shows the added dimension. The value in this dimension does not affect the value in the original dimension, so the position in this new space *C* is on a vertical line passing through *A*.

The vividness in the original solid is *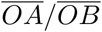* and becomes *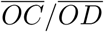*.

We can then use a long established geometric fact (it is proposition 2 in Book VI of Euclid’s Elements) that dividing two edges of a triangle by the same ratio gives points that lie on a line parallel with the third edge (and vice-versa). The line through *B* and parallel to the one passing though *C* and *A* is vertical by virtue of this proposition. This line contains point *E*, which is the position that *D* would need to be in for equal vividness. If *D* is closer to *O* than *E* then the vividness is greater.

As *B* is a boundary point, the convexity of the colour solid means that there are no points to be found further to the right on this diagram, demonstrating the proposition. In summary

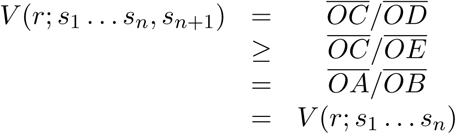

#### SI 1.2.2 Convexity of Mixing

The proposition to be demonstrated is that

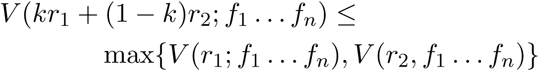

This can be be interpreted in terms of the following diagram of a plane (usually *the* plane) containing the colours of *r*_1_ and *r*_2_ and the centre of the solid:

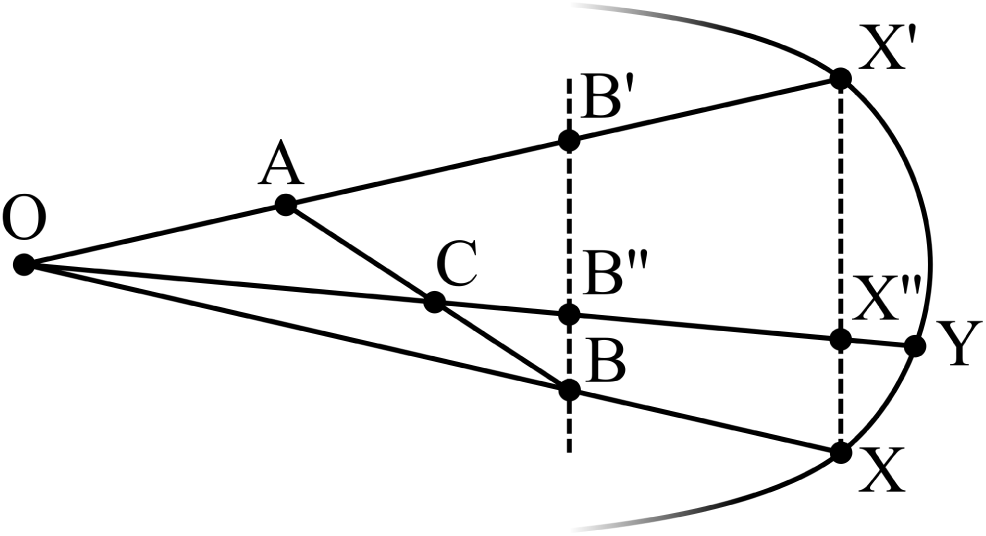

*O* is the centre of the solid. *A* and *B* correspond to the reflectance spectra *r*_1_ and *r*_2_ with the diagram drawn where *B* is the more vivid of the two. *C* is some convex combination of *A* and *B*, i.e. a point on the line between them. From the same rule used in the previous section (*Elements* VI, 2) we know that 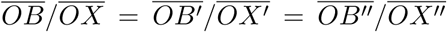. The ratio 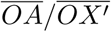 is smaller than 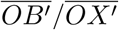 as we have chosen *B* to be the more vivid of the two.

The diagram shows that as 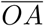 is (weakly) shorter than 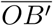, then 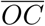 is (weakly) shorter than 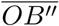 Furthermore, from the convexity of the colour solid it follows that 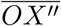 is weakly shorter than 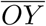. So, we can say that 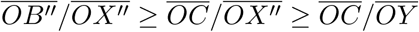. Which is the proposition to be demonstrated as 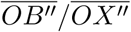 is max*{V* (*r*_1_; *x*_1_ *… x*_*n*_), *V* (*r*_2_, *x*_1_ *… x*_*n*_)*}* and 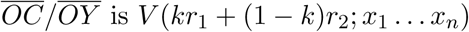. ▪

#### SI 1.3 Adjustment of Mimimal and Maximal Reflectance

Throughout this paper we have assumed that the minimal and maximal bounds on reflectance are 0 and 1. We can however work with any minimal or maximal bounds and still have a colour solid.

The colour solid is defined as the set of all colours formed by applying equation 3 to all reflectances *r* that lie between 0 and 1, we will show that if we replace the condition

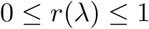

with generalised bounds

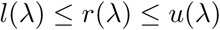

then we can construct another solid, described by

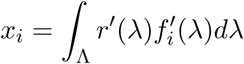

where each *x*_*i*_ is scaled between 0 and 1 and where the function *r′* lies between 0 and 1.

Let *l*(*λ*) be the lower bound of reflectance at each wavelength, and *u*(*λ*) be the upper bound. Also, let *r′*(*λ*) be a linearly scaled reflectance relative to those bounds, i.e when *r′*(*λ*) is 0, the actual reflectance is the lowest possible *r*(*λ*) = *l*(*λ*) and when *r′*(*λ*) is 1 the actual reflectance is maximal, *r*(*λ*) = *u*(*λ*). So that

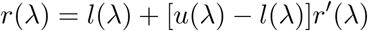

and the corresponding quantum yield is

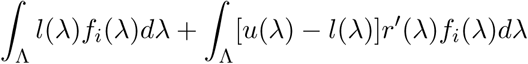

The first term is a constant offset, so may be ignored, and the second term can be seen as a rescaling of the relative yields, mathematically the same as a change in the illuminant. We can move the bounds into the relative yields and renormalise giving the relative yield functions for a new colour solid

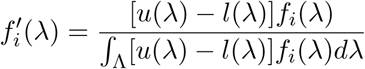

or equally, working with the illumant *l*(*λ*) and spectral sensitivities *s*_*i*_(*λ*),

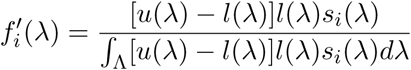

### SI 2 Numerical Solutions

#### SI 2.1 Vividness

The most difficult part of calculating vividness is finding the distance from the centre of the solid to the boundary in the direction of the chosen colour. There are two ways that this might be easily computed: One way is to calculate the geometry of the solid’s boundary, and then search the geometry for the point where a line extending from the centre through the colour crosses the boundary. This process can be optimised somewhat, but involves a lot of precalculation. Or, instead, one could use a linear programming method. This does not require precalculation, and is the method we recommend.

#### SI 2.1.1 Calculating Vividness Using Linear Programming

Finding the spectrum that lies on the boundary in a given direction from the centre can be understood as a linear programming problem. Linear programming is the name given to the optimisation of problems that have a linear objective function and linear constraints. The canonical form used for finding numerical solutions^5^ of such problems is:

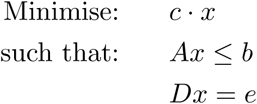

Where *x* is the vector to be determined, *b* and *c* are vector, and *A* matrix, parameters. Our task is to put the problem of finding the distance to the boundary into a form that can be understood by standard linear programming routines, such as the simplex method.

In this problem, we find the boundary colour by first finding the spectrum which lies on the boundary of solid. We must work with spectra, not colours, as the colour solid is determined by a constraint on the form of the spectrum.

**Inequality Constraints** (form of *Ax ≤ b*)

If *x* is a vector representing a reflection spectrum, then we have constraints of the form

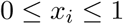

which we can write in block matrix form as:

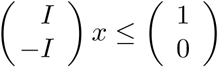

**Equality Constraints** (form of *Dx* = *e*)

There are also equality constraints. The colour of the boundary spectrum must lie on a line passing through the reference colour and the centre. The colours (*s*) lying on this line are given by:

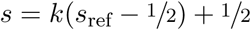

where *s*_ref_ is the reference colour and *k* is a variable giving the position along the line. ½ is used here to represent the vector (½, ½ *…*).

This has the form *s* = *kp*+*q*, where *p* = *s*_ref_ − ½ and *q* = (½, ½ *…*) are vectors. To get this in the form of *Dx* = *e* that is needed to specify the linear programming problem, we need to do a little algebraic manipulation. If can we write the constraint of the colour to a line in the standard form: *Mc* = *v*, then it is simple to see that when *F* is a matrix containing the discretised relative quantum yield functions (see equation eqn:discrete), the colour *s* is given by *Fx* and we have *MFx* = *v*, meaning that *D* = *MF* and *e* = *v*.

There are multiple choices of *M* and *v* which constrain the colour in exactly the same way. One way of obtaining an *M* and *v* is by picking one dimension for which *p* is non-zero (the ‘pivot’), using this to solve for *k* and substituting the solution back. The result of this procedure is an *n*-by-*n* − 1 matrix and an *n* − 1 vector. The matrix has can be understood as having ones on the leading diagonal and −*p*_*i*_*/p*_pivot_ in the pivot column, with the pivot row removed. For example, for a tetrachromat pivoting with *i* = 2 we have, the 4-by-3 matrix:

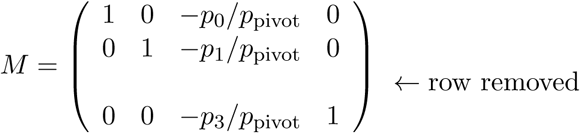

The corresponding vector has values of the form *q*_*i*_ − (*q*_pivot_*/p*_pivot_)*p*_*i*_ and, like *M*, has the pivot row removed.

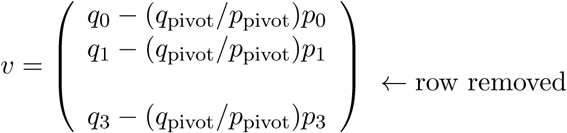

**Objective Function** (form of *c · x*)

Lastly, we need the objective function which is to be maximised. This measures how far we are from the centre of the solid. A dot product using colour is the choice because it is signed (unlike the Euclidean distance), and is linear. The function

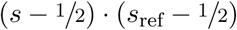

is the distance from the centre, projected in the direction of *c*_*r*_*ef*. To get this in the form *c · x* we rewrite it as:

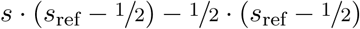

The second term is a constant. This means that its value does not effect the minimisation and we can ignore it, giving:

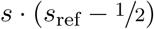

After substituting *Fx* for *s*, and rearranging using the commutativity of the dot product we find that

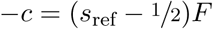

The minus sign is present because the problem is specified as a minimisation, not a maximisation.

#### SI 2.2 Colour Solid Geometry

The geometry of the colour solids have been calculated here using a convex pruning method.

Through convexity it is possible to demonstrate that the boundary of the colour solid is formed by reflectances that contain only zeros and ones. Every spectrum can be formed though a convex combination of zero-one spectra, thus any colour must be a convex combination of the colours formed by zero-one spectra.

To calculate the solid numerically, we use discretised relative quantum yield functions, *g*_*i*_(*j*), for a set of wavelengths *λ*_0_, *λ*_1_, *… λ*_*j*_ *… λ*_*n*_.

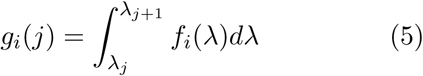

where *i* indexes the photoreceptor class, and *j* indexes the wavelength band. The algorithm works by incrementally enumerating all the reflectance spectra that may lie on the surface of the solid, ignoring all those for which this is clearly impossible.

Denote the set of colours defined by non-zero discretised reflectance only in the region [*λ*_0_, *λ*_*k*_] by 𝒮_*k*_. This will be uniquely defined by a finite set of points lying on the convex hull of that set, *ℋ*_*k*_. The sets *S*_*k*_ and *S*_*k-*1_ are related to each other by:

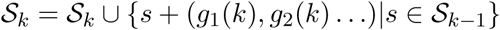

The first and second terms of the right hand sum correspond to the addition of reflectances in region [*λ*_*k-*1_, *λ*_*k*_] with values 0 and 1 respectively. Thus *S*_*n*_ contains all colours corresponding to discretised reflectances with zero or one everywhere, and has 2^*n*^ elements. For resolutions as low as *n* = 10 the full enumeration of this can be problematic. Even more so when it comes to obtaining the convex hull, the algorithms for which which are super-linear in both the number of points and the dimension. To avoid having such large sets we prune away fruitless candidates as we go along.

For calculating the bounding points of the 𝒮_*k*_, only the values *ℋ*_*k* − 1_ matter. This is because the addition of a vector, *v*, to any point in 𝒮_*k*_ will result in a point that is a convex combination of points in *{s* + *v|s ∈ ℋ*_*k*_*}*. If we let *𝒞* denote the operation which obtains the points defining the convex hull, then it follows that:

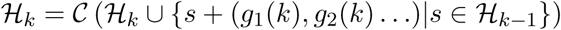

Which, when applied iteratively, results in a computationally feasible number of points. The speed of this procedure may be increased by including the results of a low resolution version of the calculation (even just the white point). Call these known points *𝒦*, and then instead iterate using:

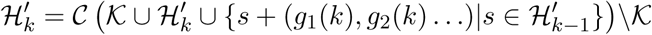

with:

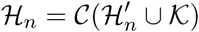

The last application of *C* may be omitted if we know that *𝒦 ⊆ ℋ*_*n*_ – a condition which is desirable for the efficiency of the pruning.

The computational complexity of this algorithm depends on the number of photoreceptor classes, and becomes difficult for observers with more that 5 photoreceptors.

#### SI 2.3 Using Linear Programming

A linear programming method like the one described in the vividness calculation can be used to obtain a collection of points describing the geometry of the colour solid. To do this, one samples directions around the centre, and calculates the corresponding boundary spectra using linear programming. This method is has a disadvantage in comparison with the method below in that it can only resolve fine detail such as points or sharp edges by using a very high density of points.

#### SI 2.4 Metameric Colours

This algorithm can then be used to calculate the colours for one observer that correspond to the colours for another. Call the discretised relative quantum yield functions for the two observers *a*_*i*_(*j*) and *b*_*i*_(*j*) respectively, and let 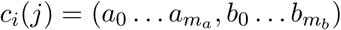 where *m*_*a*_ and *m*_*b*_ is the number of photoreceptor classes for *a* and *b*. An *m*_*a*_ +*m*_*b*_ dimensional colour solid can then be constructed for *c*_*i*_. The set of observer *b* colours corresponding to an observer *a* colour can then be calculated as the *m*_*b*_ dimensional cross-section of the *m*_*a*_ + *m*_*b*_ dimensional solid for which the observer *a* coordinates are constant.

As this involves calculating a *m*_*a*_ + *m*_*b*_ dimensional colour solid, it is only ‘fast’ for *m*_*a*_ +*m*_*b*_ *<* 6. For other cases, a dynamic programming approach is prudent.

### SI 3 Purity in Multiple Dimensions

If colour purity – the saturation relative to the maximally saturated colour of a given hue – is generalised to arbitrary dimension one obtains a quantitiy very similar to vividness. Much as with the extreme spectra of the object colour solid, the border of an arbitrary chromaticity diagram is given by the linear combinations of a few spectra, and as in the colour solid, when the dimension increases, the new boundary will include spectra from the lower dimensional boundary. These spectra are monochromatic, and play the same role as the extreme spectra. But the kind of results that purity yields cannot so easily be employed in developing the kinds of arguments that we developed in the main text, even in approximation.

One difficulty lies in the spectra of the purest colours being monochromatic (or combinations of a few monochromatic spectra). Monochromatic lights are particularly unnatural; an approximately monochromatic *reflectance* has the property of reflecting less and less light as it becomes more pure – reflecting none at the point of perfect monochromaticity.

Another difficulty is that observers with different numbers of photoreceptor classes (different dimensionality) have boundary sets that differ significantly in quality.

For monochromats there is no useful definition of saturation, and therefore no saturation based vividness-like measure exists.

In all other cases, the neatest formulation of the boundary is as the convex hull of all the monochromatic lights. If the monochromatic lights form a convex figure, the convex parts will not be part of the boundary. This means that for dichromats the chromaticity space is bounded by monochromatic two lights, one positioned at the wavelength with the highest ratio in sensitivities and one at the lowest *λ*_low_ and *λ*_high_:

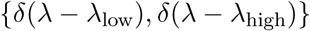

Unlike the dichromat Schrödinger spectra, these are completely dependent on the spectral sensitivities of the photoreceptors.

For trichromats the boundary often contains a good portion of the monochromatic locus and there is often only one linear connecting sections spanning from a short wavelength to a long one: the ‘purple line’:

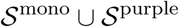

The dimensionality of the set of monochromatic lights, and of the purple line is the same.

With tetrachromacy and above, the chromatic boundary becomes dominated by non-spectral (i.e. not monochromatic) spectra, most of the boundary is formed by a *n* − 2 dimensional surface filling in between the the 1 dimensional locus of monochromatic spectra. The surface is formed by linear combinations of *n* − 2-tuples of monochromatic lights, and typically there will be a small number of dominant monochromatic light that are part of the combinations covering large portions of the surface – in other words, often, a bit of the monochromatic locus “sticks out”. Determining which bits stick out is not particularly simple, and whilst there are rules of thumb that help, it makes framing the arguments we have made here much more complicated. This is aside from the fact that the object colour solid is simply the most appropriate model for reflectance.

## Footnotes

1 The maximal reflectance could instead be set according to a standard such as BaSO_4_ pellet – this does not affect the arguments we make here (see appendix SI 1.3) and should, in fact, result in a more practical measure of vividness.

2 When discretised approximations like those in the appendix are used to calculate boundary spectra, a step transition between two discretisation points will appear at an intermediate intensity. This is a discretisation artifact.

3 For a fully formal approach we require another rule stating that *V* is not dependent on the order in which the relative quantum yield functions are specified.

4 Usually there is only one spectrum corresponding to a point on the boundary of the solid, but there might be more in cases where the surface is perfectly flat. Inside the boundary, or at points on the boundary without non-zero bounded curvature, there is an infinite number. Measuring the absolute sizes of the sets of spectra corresponding to a single colour is non-trivial because there is no ‘infinite-dimensional’ analogue to the Lebesgue measure. In this work we avoid this problem by always comparing between two different finite dimensional colour solids (*⊆* [0, 1]^*n*^), rather than looking for a way of measuring spectrum-space ([0, 1]^*∞*^).

5 The equality constraints (*Dx* = *e*) are usually explicit in numerical procedures, but in mathematical treatments they are usually understood as a pair of inequality constraints: *Dx ≤ e* and *−Dx ≤ −e*.

